# Discovery and Functional analysis of a novel *ALPK1* variant in ROSAH syndrome

**DOI:** 10.1101/2024.09.13.612837

**Authors:** Tom Snelling, Leo O. Garnotel, Isabelle Jeru, Maud Tusseau, Laurence Cuisset, Antoinette Perlat, Geoffrey Minard, Thibaut Benquey, Yann Maucourant, Nicola T. Wood, Philip Cohen, Alban Ziegler

## Abstract

ROSAH syndrome is an autosomal dominant autoinflammatory disorder characterised by visual disturbance caused by pathogenic variation in the protein kinase ALPK1. Only two such variants have been reported to cause ROSAH syndrome to date: 66 out of 67 patients harbour the Thr237Met variant, while a single patient carries a Tyr254Cys variant. Here we identify a family in which ROSAH syndrome is caused by a Ser277Phe variant in *ALPK1*. The phenotypic variability in this family is high, with four of the seven individuals legally blind. Hypohidrosis, splenomegaly and arthritis were present in several family members. In contrast to wildtype ALPK1, which is activated specifically by the bacterial metabolite ADP-heptose during bacterial infection, ALPK1[Ser277Phe] was also activated by the human metabolites UDP-mannose and ADP-ribose, even more strongly than the ALPK1[Thr237Met] variant. However, unlike ALPK1[Thr237Met], ALPK1[Ser277Phe] could additionally be activated by GDP-mannose. These observations can explain why these *ALPK1* variants are active in cells in the absence of ADP-heptose and hence why patients have episodes of autoinflammation. Examination of the three-dimensional structure of ALPK1 revealed that the sidechains of Ser277 and Tyr254 interact but mutational analysis established that this interaction is not critical for the integrity of the ADP-heptose binding site. Instead, it is replacement of Ser277 by a large hydrophobic phenylalanine residue or the replacement of Tyr254 by a much smaller cysteine residue that is responsible for altering the specificity of the ADP-heptose-binding pocket. The characterisation of *ALPK1* variants that cause ROSAH syndrome suggests ways in which drugs that inhibit these disease-causing variants selectively can be developed.

## Introduction

ALPK1 (Alpha-protein kinase 1) is a key component of an innate immune signalling pathway that is activated by the bacterial nucleotide sugars ADP-D-glycero-β-D-manno-heptose (ADP-D,D-heptose) and ADP-L,D-heptose (hereafter ADP-heptose) [1]. The binding of either of these ADP-heptoses within the N-terminal domain of ALPK1 activates the C-terminal catalytic kinase domain, enabling ALPK1 to phosphorylate TIFA (TRAF-interacting protein with forkhead-associated domain) [1]. This leads to the polymerisation of TIFA and the formation of a signalling complex that activates the transcription factors NF-κB (nuclear factor kappa-light-chain-enhancer of activated B cells) and AP-1 (activator protein 1) [1–4].

ROSAH (<u>R</u>etinal dystrophy, <u>O</u>ptic nerve oedema, <u>S</u>plenomegaly, <u>A</u>nhidrosis and migraine <u>H</u>eadache) syndrome is an autosomal dominant disease caused by pathogenic variants in *ALPK1* [5]. Patients with ROSAH syndrome usually present in the clinic with failing eyesight but, beyond this, the precise phenotype varies and can additionally include autoinflammatory conditions, such as arthritis [6]. In all but one patient with ROSAH so far identified, the variant Thr237Met of ALPK1 was found, the exception being a single patient with a Tyr254Cys variant [7]. Thr237 and Tyr254 are located within the N-terminal ADP-heptose binding domain of ALPK1, but only Thr237 forms a direct interaction with ADP-heptose itself [1].

The overexpression of ALPK1[Thr237Met] or ALPK1[Tyr254Cys] in HEK293 (human embryonic kidney 293) cells activates NF-κB/AP-1-dependent gene transcription in the absence of ADP-heptose in the cell culture medium, implying that these mutated proteins are activated by an ADP-heptose-independent mechanism in cells [4]. However, when immunoprecipitated from cell extracts and assayed for kinase activity, ALPK1[Thr237Met] is devoid of kinase activity in the absence of ADP-heptose, similar to the wild-type (WT) enzyme [4]. These observations led to the unexpected finding that, in contrast to WT ALPK1, ALPK1[Thr237Met] can also be activated by the human nucleotide sugars UDP-α-D-mannose and ADP-D-ribose in cell free assays [4]. This suggested that activation of ALPK1[Thr237Met] by one or more human sugar nucleotides may cause chronic activation of ALPK1 in cells and underlie autoinflammation in ROSAH syndrome. In contrast to ALPK1[Thr237Met], the ALPK1[Tyr254Cys] mutant was devoid of activity when assayed in this way, even in the presence of ADP-heptose, indicating that it is unstable when removed from cells.

So far, 67 patients with ROSAH syndrome from 29 unrelated families and carrying only two different missense variants have been reported [5–10]. However, as the first case was only described in 2019, the condition is probably underdiagnosed and further patients with the disease are likely to be identified. Here we report on the identification of an additional family with ROSAH syndrome caused by a new Ser277Phe variant in *ALPK1* and characterise its effect on ALPK1 activity.

## Materials and Methods

### Genetic studies

Diagnostic laboratories performed genetic analyses on genomic blood DNA extracted from peripheral blood leukocytes using standard procedures. Genome sequencing for individuals II1 and III1 was performed at the SeqOIA laboratory (LBMS SeqOIA, Paris, France). FASTQ files were obtained from the bcl2fastq demultiplexing tool (v2.20.0.422, Illumina) and aligned to the GRCh38.92 genomic reference using BWA-MEM (v0.7.15). Haplotype Caller (v4.1.7.0) was used to call Single Nucleotide Variants (SNVs) and delins (<50bp); variants were annotated by SNPEff (v4.3t). Structural variants were detected by ClinSV (v1.0.1) and WiseCondor (v1.2.4). The resulting SV were then annotated with AnnotSV (v3.0.7) [11]. In-house software (GLEAVES, v3.3.23) was used for the variant interpretation and reporting. Further details are available on the website of the LBMS SeqOIA platform (https://laboratoire-seqoia.fr/). Exome sequencing for individual II2 was performed at Biomnis laboratory (Lyon, France) using a Human Exome 2.0 Plus Comprehensive Exome library and a Novaseq 6000 sequencer.

### Antibodies

Antibodies recognising GAPDH (#2118) and anti-rabbit IgG (#7074) were from Cell Signalling Technology and an antibody recognising FLAG (#F3165) was from Sigma-Aldrich.

### DNA constructs

The following DNA plasmids encoding FLAG-tagged WT or variant forms of ALPK1 with expression under the control of the CMV promoter were made by Medical Research Council Reagents and Services, Medical Research Council Protein Phosphorylation and Ubiquitylation Unit, University of Dundee, and assigned unique identifiers (listed in parentheses); these constructs are available on request (mrcppureagents.dundee.ac.uk): WT (DU65668), R150A (DU71740), R150A/T237M (DU71743), R150A/Y254C (DU71741), R150A/S277F (DU71954), T237A (DU71986), T237M (DU65723), Y254C (DU71685), Y254F (DU71987), S277A (DU71988), S277F (DU71952) and S277F/K1067M (DU71956).

### Nucleotide sugars

The sources of nucleotide sugars are as described (Snelling et al 2023), except that the ADP-heptose used in this study was ADP-L,D-heptose (#tlrl-adph-l) from Invivogen. All nucleotide sugars were prepared as 1 mM stock solutions.

### Transfection of HEK293-Blue cells in 96-well format and measurement of NF-κB/AP-1 gene transcription

Cells (60,000) in 0.1 ml of Dulbecco’s modified Eagle’s medium containing 10% (v/v) heat-inactivated foetal bovine serum were “reverse” transfected in a 96-well plate using 1 µl of lipofectamine 2000 (ThermoFisher, #11668019) and 0.4 µg of plasmid DNA. After 24 h the culture medium was aspirated and replaced by 75 µl of Dulbecco’s modified Eagle’s medium containing 10% (v/v) heat-inactivated foetal bovine serum, 100 U/ml penicillin and 0.1 mg/ml streptomycin with or without 5 μM ADP-heptose. This culture medium was then collected after a further 24 h, and 30 µl was incubated with 170 µl of QUANTI-blue solution (Invivogen, #rep-qbs) in a new 96-well plate. The plate was incubated for 30 min at room temperature and the absorbance at 645 nm measured using a microplate reader. For detection of ALPK1 expression by immunoblotting, the cells were lysed in 30 µl sodium dodecyl sulphate (SDS) sample buffer (Millipore, #70607) supplemented with 0.5% (v/v) benzonase endonuclease (Sigma-Aldrich, #E1014), 1 mM MgCl2 and protease inhibitor cocktail (Roche, #11836170001).

### Cell-free ALPK1 phosphorylation assays

Plasmid DNA (60 µg) encoding WT or mutant ALPK1 was transfected into 15 cm dishes of ALPK1 KO HEK293-Blue cells using 150 µl of lipofectamine 2000. After 24 h, the cells were washed twice with PBS and scraped in 1 ml ice-cold lysis buffer containing 50 mM Tris–HCl pH 7.5, 1 mM EDTA, 1 mM EGTA, 1% (v/v) Triton X-100, 2 mM dithiothreitol (DTT), 270 mM sucrose supplemented with protease inhibitor cocktail (Roche, #11836170001). Cell lysates were clarified by centrifugation for 20 min at 20,000 x g at 4°C and the supernatants (cell extracts) transferred to 1.5 ml microcentrifuge tubes. Cell extract protein (0.05 mg) containing WT FLAG-ALPK1 or FLAG-ALPK1 mutants (normalised for ALPK1 expression) were incubated for 1 h at 4°C on a rotating wheel with 15 μl of packed anti-FLAG M2 affinity gel (Sigma-Aldrich, #A2220), which had been washed twice with lysis buffer prior to use. After centrifugation for 30 sec at 1000 x g at 4°C, the supernatant was discarded and the pelleted gel washed three times with 50 mM Tris-HCl pH 7.5, 2 mM DTT, 0.1% Triton X-100 (Buffer A) containing 500 mM NaCl, twice with Buffer A and once with 50 mM Tris-HCl (pH 7.5), 2 mM DTT, 0.1 mM EGTA and 10 mM magnesium acetate (Buffer B). The immunoprecipitated FLAG-ALPK1 was then incubated at 30 °C in 25 µl of Buffer B containing 2.1 μM glutathionine-S-transferase (GST)-TIFA (dialysed against 50 mM Tris-HCl pH 7.5, 2 mM DTT) and 1 mM [γ-^32^P]ATP (specific radioactivity 500 cpm/pmol). The reactions were initiated by the addition of the [γ-^32^P]ATP and terminated after 30 min by the addition of lithium dodecyl sulphate sample buffer containing 2.5% (v/v) 2-mercaptoethanol and heated for 5 min at 75°C. The FLAG resin was pelleted by centrifugation for 30 sec at 13,000 x g and the supernatant subjected to SDS-PAGE. After staining for 30 min with InstantBlue Protein Stain (Abcam, #ab119211) and destaining for 16 h in water with frequent changes, the bands corresponding to GST-TIFA were excised and the incorporation of ^32^P-radioactivity analysed by Cerenkov counting.

**Other procedures** All other methods, including the origin and culturing of cells, have been described elsewhere [4].

## Results and Discussion

### Clinical Investigation of a family harbouring the symptoms of ROSAH syndrome

Individual I1 is the mother of individuals II1 and II2 (**Fig 1A**). She died from breast cancer but had a history of progressive visual impairment from her 20s and legal blindness in her 40s. However, her DNA was not available for sequencing.

**Figure 1:**
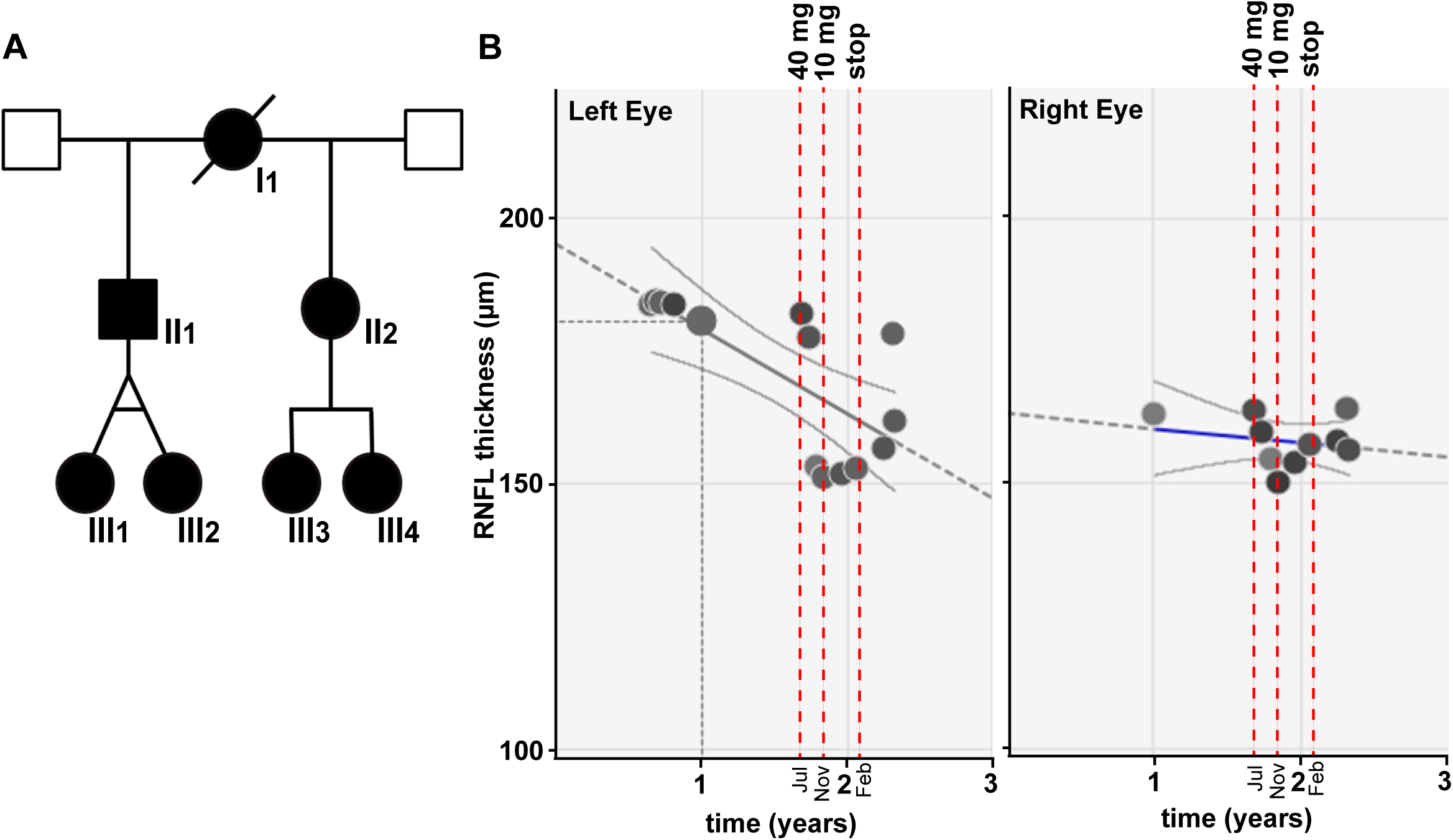
Family expressing the ALPK1[Ser277Phe] variant and response of Individual II2 to systemic corticosteroid therapy. **(A)** Family Tree. Individual II1 and individual II2 have the same mother but different fathers. The daughters of individual II1 are monozygous twins. (**B**) Retinal Nerve Fibre Layer (RNFL) thickening showed an initial improvement with systemic corticosteroid therapy (40 mg) but when treatment was reduced to 10 mg and then stopped, the RNFL thickness returned to the level observed prior to corticosteroid therapy.

Individual II1, a 49-year-old man is the father of individuals III1 and III2 and the maternal half-brother of individual II2 (**Fig 1A**). He has been followed since childhood for visual impairment, with an intermediate and posterior uveitis, associating hyalitis, and bilateral papilloedema, major macular oedema and preretinal neovessels. He was recognised legally blind in his 40s. He also had painful arthralgia of the hands and knees. He reported hypohidrosis but had no history of recurrent headaches or haematuria.

Individual II2 (**Fig 1A**) is a 38-year-old woman who presented with bilateral eye pain and visual loss. Personal medical history was marked by unclear episodes of recurrent ocular inflammation since the age of 20. She reported numerous episodes of macroscopic haematuria, most often associated with fever. Arthralgia of the hands consecutive to a deforming arthritis, hypohydrosis, splenomegaly, multiple caries, and parotidomegaly were also found upon examination. No recurrent headaches were reported.

Initial best corrected visual acuity of individual II2 was 20/32 in both eyes. Intra-ocular pressure was normal (15 mm Hg on the right-eye and 16 mm Hg on the left eye). Slit-lamp examination found a bilateral inflammation in the anterior chamber of the eyes with granulomatous central keratic precipitates. A 1^+^ vitritis associated with inferior snowballs was seen bilaterally. No retinal or choroidal foci were found, and nor was retinal vasculitis. However, a deep vascular attenuation was seen from mid- to extreme periphery which was associated with ghost retinal vessels. A mild optic disc oedema was present on both eyes.

Autofluorescence showed patterns of hyperfluorescence predominantly in temporal zones and juxta-vascular zones. A whole posterior pole leakage was found on fluorescein angiography, mostly seen around the optic disc and the juxta-vascular areas. Infracyanine green angiography was considered normal. A bilateral retinal nerve fibre layer (RNFL) thickening was confirmed on optical coherence tomography (OCT), and macular cysts in inner retinal layers were found on macular OCT (**Supplementary Fig 1** and **Supplementary Fig 2**). Routinely conducted laboratory tests for uveitis were negative.

Initial management with topical corticosteroid drops resolved anterior inflammation, with no effect on posterior signs. She was lost on follow up and presented a year after with a recurrence of bilateral anterior inflammation. Systemic corticoids (1 mg/kg) were then introduced and tapered with a good efficacy on macular and optic disc thickening up to 10 mg (**Fig 1B**). Given this partial response, an anti-IL-6 therapy (tocilizumab) was introduced to further reduce inflammation and thickening of the posterior retina and optic disc. A follow-up, 4 months after the initiation of this treatment highlighted decreased macular and optic disc thickening and improved visual acuity.

Individuals III1 and III2 (**Fig 1A**) are 20-year-old female monozygotic twins. They both present with identical symptoms. They were born prematurely at 34 weeks of gestation and have had delays in learning and speech development. Visual acuity at the age of 10 was normal. At the age of 12, during an evaluation for decreased visual acuity, a diagnosis of bilateral macular and papillary oedema with preretinal neovascularization was made, along with a clinical picture of polyarthralgia and finger deformities characterized by broadening of the proximal interphalangeal joints without active synovitis. At the age of 14, the visual acuity of individual III1 was 1.2/10 P12 for the right eye and 1.4/10 P12 for the left eye; that of individual III2 was 9/10 P2 for the right eye and 0.8/10 P28 for the left eye. They also have autoimmune hypothyroidism, which was diagnosed at the age of 12 in both patients.

The two patients were initially treated for the macular oedema with systemic corticosteroids and local steroid injections. Systemic corticosteroids did not reduce the macular oedema, but iterative local injections of corticosteroids were successful. Then, to reduce corticosteroid use, methotrexate was introduced, followed by adalimumab and then infliximab. Intravenous tocilizumab was started in 2021, which was effective in combination with local dexamethasone injections, transitioning to subcutaneous injection for individual III2. Individual III2 developed cutaneous allergy to tocilizumab, necessitating a switch to a JAK kinase inhibitor (baricitinib). In addition to this systemic treatment, injections of fluocinolone acetonide implant for individual III1. and dexamethasone implants for individual III2 are performed once or twice a year. Multiple dexamethasone injections triggered bilateral cataracts in both patients, which were operated on between 2020 and 2022.

Pre-retinal haemorrhages associated with the neovessels were also noted, for which a few laser sessions and intravitreal anti-VEGF injections were performed. The last hemorrhage was identified at the age of 16 in the 2 patients, with no recurrence since.

The latest acuities at the age of 21 were: 2/10th P14 in both eyes for individual III1 and 1.6/10ths P8 on the right eye and 1.4/10ths P16 on the left eye for individual III2. Intraocular pressures have always been normal, and they did not present uveitis.

Individuals III3 and III4 are the daughters of individual II2 (**Fig 1A**). At the last evaluation at age 11, III3 had a normal fundus, but the macular and RNFL OCTs showed a discrete thickening of RNFL and internal retinal layers. Individual III4 at the last evaluation was age 7 and an optic nerve head oedema was clearly visible on the fundus. Her macular and RNFL OCTs also showed a thickening of RFNL and internal retinal layers. In both cases, oedema was pre-eminent along the path of RFNL and retinal vasculature. As visual function was not impaired and clinical inflammation was low, no treatment has yet been undertaken for them.

The phenotypic variability in this family is high, especially concerning the severity of the visual impairment with 4 individuals (I1, II1, III1 and III2) legally blind while the other three (II2, III3, and III4) have none to mild visual impairment. Notably, this variability is not only related to the age as individual II2 with a mild visual impairment is older than her nieces (individuals III1 and III2). It could be hypothesized that this less severe phenotype is related to a genetic contribution from the father of individual II2.

As the diagnosis has been made at a young age for individuals III3 and III4 before the onset of visual loss, we will follow them twice a year to determine whether a treatment should be introduced. This follow-up should improve the understanding of the natural history of the disease and whether some clinical features, such as the intensity of changes on fundus examination, correlates with the likelihood of vision loss. This information will be critical to identify surrogate markers of efficacy for future clinical trials.

### Characterisation of a novel *ALPK1* variant

Genome sequencing was performed in subjects II1 and III1 and exome sequencing in subject II2. This led to the identification in these affected members of a heterozygous missense variant in exon 10 of *ALPK1*: c.830C>T; p.Ser277Phe (transcript NM_0251144.4). The segregation of the variant was confirmed by Sanger sequencing in subjects III2, III3 and III4. Notably, we did not identify any alternative molecular etiology compatible with the disease phenotype in any of these three patients. Several additional lines of evidence supported the causal role of this variant in the disease phenotype. The variant was absent from databases reporting variants from the general population (gnomAD v4.1.0), as well as from ClinVar, a database that aggregates information about genomic variations and their relationship to human diseases. The p.Ser277Phe variant identified herein affected an amino acid strongly conserved in all major vertebrate lineages. Notably, the variant pathogenicity predictions made by the bioinformatic tools CADD Phred: 24.3; Mistic: 0.36; REVEL: 0.229; AlphaMissense: 0.426 remained uncertain. Similar observations were made when using the same bioinformatic tools with the other *ALPK1* pathogenic variants published previously (p.Thr237Met, p.Tyr254Cys). Indeed, this type of tool performs better in predicting deleterious effects for loss-of-function variants than for gain-of-function variants [12]. According to the international recommendations for variant classification [13], the variant identified could not be classified as “pathogenic” in the absence of additional clues, especially functional assays showing a deleterious effect. This underscores the need for cell-based tests to assess the functional impact of the p.Ser277Phe variant on ALPK1 activity and hence the additional experiments described below were carried out.

### Effect of the ALPK1[Ser277Phe] mutation on NF-κB/AP-1-dependent gene transcription in HEK293 cells

We initially compared gene transcription induced by ALPK1 mutants after re-expressing them in ALPK1 knockout (KO) HEK293 cells, which contain a synthetic gene encoding secreted embryonic alkaline phosphatase under the control of NF-κB and AP-1 promoters. In this assay, we found that gene transcription induced by ADP-heptose was similar in cells expressing WT ALPK1, ALPK1[Ser277Phe], ALPK1[Thr237Met] or ALPK1[Tyr254Cys] (**Fig 2A**). When ADP-heptose was omitted, gene transcription was negligible in cells expressing WT ALPK1, but was still observed in cells expressing ALPK1[Ser277Phe], ALPK1[Thr237Met] or ALPK1[Tyr254Cys] (**Fig 2A**). The extent of gene transcription induced by ALPK1[Ser277Phe] and ALPK1[Tyr254Cys] was greater than that induced by the ALPK1[Thr237Met] mutant (**Fig 2A**). The level of expression of the ALPK1[Ser277Phe] mutant in transfected ALPK1 KO cells was similar to WT ALPK1 (**Supplementary Fig 3**).

**Figure 2:**
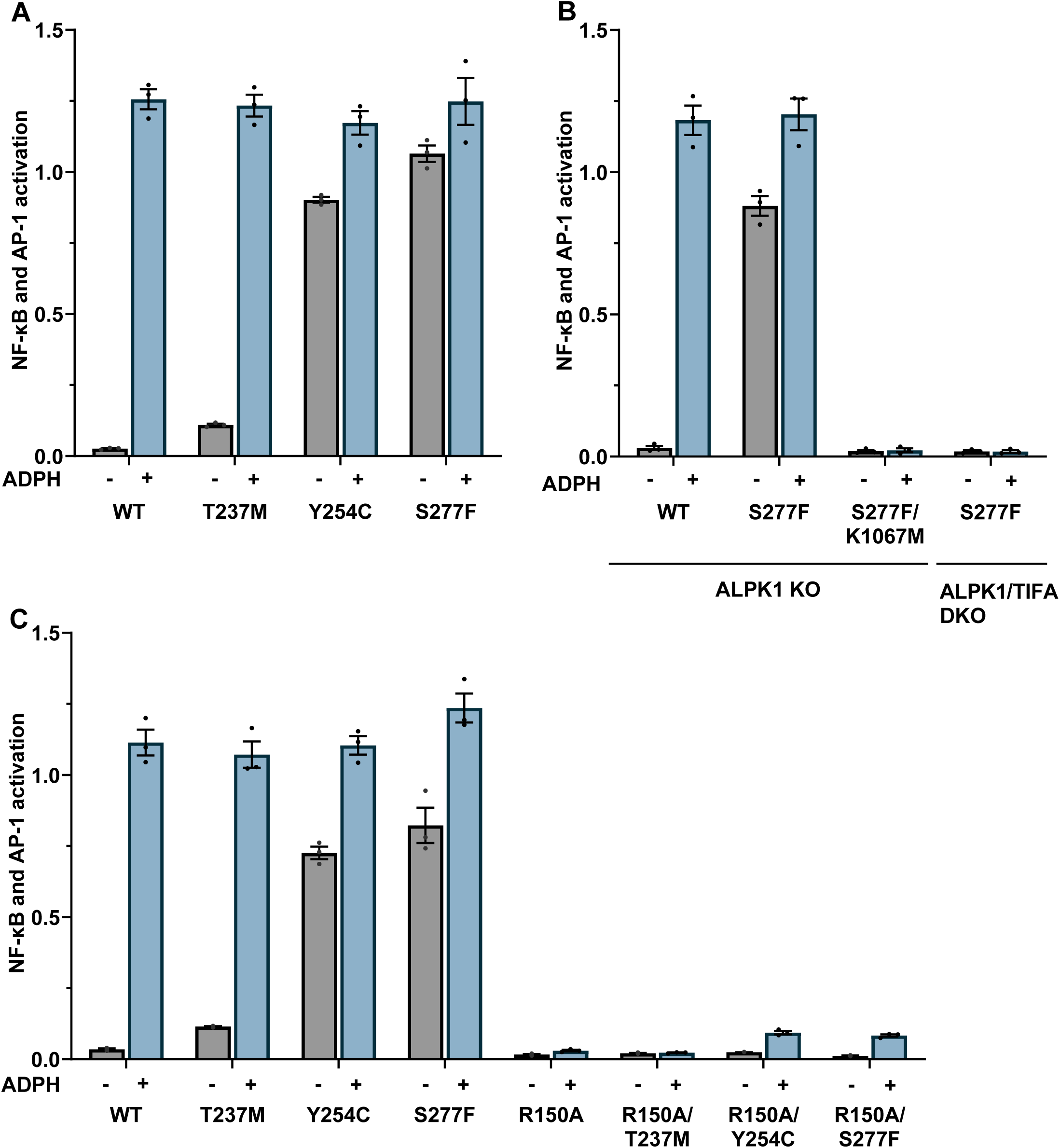
The ALPK1[Ser277Phe] mutant stimulates NF-κB/AP-1-dependent gene transcription in the absence of ADP-heptose. **(A)** ALPK1 KO cells were transfected with plasmids encoding WT ALPK1 or the indicated ALPK1 mutants. 24 h later, cells were incubated with (blue bars) or without (grey bars) 5 μM ADP-heptose (ADPH) and NF-κB/AP-1-dependent gene transcription was measured after another 24 h (see Methods). **(B)** ALPK1 KO or ALPK1/TIFA double KO (DKO) cells were transfected with the plasmids indicated and analysed as in (A). **(C)** As in (A) but using plasmids in which each ROSAH-causing mutant was combined with the mutation Arg150Ala, a residue that is critical for ADP-heptose binding. **(A-C)** Results are shown as the mean with error bars indicating SEM from an experiment performed in triplicate. Similar results were obtained in two additional experiments.

The activation of NF-κB/AP-1-dependent gene transcription by ALPK1[Ser277Phe] did not occur in TIFA KO cells, or when the Ser277Phe mutation was combined with the Lys1067Met mutation (**Fig 2B**). The latter mutation is located within the kinase domain and disrupts ATP binding at the catalytic site [1]. Thus, the kinase activity of ALPK1 and the expression of TIFA are both required for ALPK1[Ser277Phe] to induce gene transcription, irrespective of whether ADP-heptose is present. These results are similar to those obtained previously with the ALPK1[Thr237Met] and ALPK1[Tyr254Cys] variants and this suggested that, like these variants [4], ALPK1[Ser277Phe] might be activated in cells by human nucleotide sugars.

To investigate if ALPK1[Ser277Phe] was active in cells because it was being activated by another molecule binding to the same site as ADP-heptose, we disrupted the ADP-heptose binding site. Arg150 is an amino acid residue that has been shown to be critical for ADP-heptose binding because it forms electrostatic interactions with the negatively charged phosphate groups of ADP-heptose within the ADP-heptose-binding site [1]. We found that mutation of Arg150 to Ala not only abolished (WT ALPK1 or ALPK1[Thr237Met]) or greatly reduced (ALPK1[Tyr254Cys] or ALPK1[Ser277Phe]) gene transcription in the presence of ADP-heptose, but also activity in the absence of this nucleotide sugar (**Fig 2C**). This was not caused by failure of the “double mutant” proteins to express, although expression was reduced relative to the “single” mutants (**Supplementary Fig 3**). The finding that the ALPK1[Arg150Ala/Ser277Phe] double mutant was unable to stimulate gene transcription supported the hypothesis that ALPK1[Ser277Phe] was being activated in human cells by another molecule that nevertheless was engaging the ADP-heptose binding site and led us to study the effects of human nucleotide sugars.

### Effect of the ALPK1[Ser277Phe] mutation on ALPK1 activity in cell-free kinase assays

To identify human metabolites that might activate ALPK1[Ser277Phe], we assayed its activity in cell-free kinase assays. We found that, similar to ALPK1[Thr237Met], ALPK1[Ser277Phe] was activated by UDP-α-D-mannose and ADP-D-ribose, but more strongly than ALPK1[Thr237Met] (**Fig 3A**). Moreover, the activation of ALPK1[Ser277Phe] by UDP-α-D-mannose and ADP-D-ribose was prevented by combination with the Arg150Ala mutation (**Fig 3B**), indicating that the mammalian nucleotide sugars activate ALPK1[Ser277Phe] by binding to the ADP-heptose binding site. Interestingly, and unlike ALPK1[Thr237Met] or WT ALPK1, ALPK1[Ser277Phe] was additionally activated by GDP-α-D-mannose (**Fig 3A**). The activity of the ALPK1[Ser277Phe] mutant in the presence of ADP-heptose was lower than that of ALPK1[Thr237Met] in the presence of ADP-heptose which, in turn, was lower than that of WT ALPK1 (**Fig 3A**).

**Figure 3:**
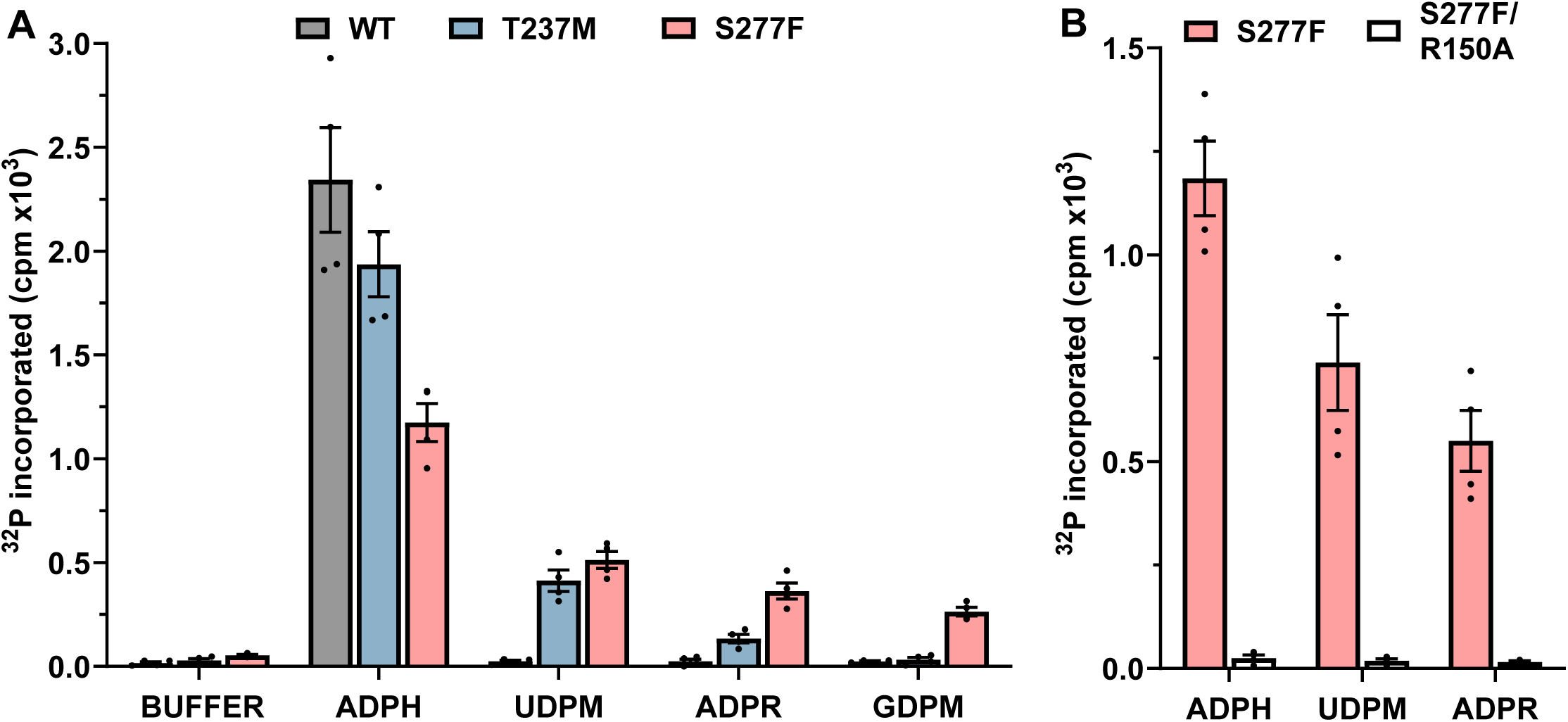
ALPK1[Ser277Phe] can be activated by mammalian nucleotide sugars in cell-free protein phosphorylation assays. **(A)** FLAG-tagged WT ALPK1 (WT, grey bars), ALPK1[Thr237Met] (T237M, blue bars) or ALPK1[Ser277Phe] (S277F, pink bars) were immunoprecipitated from cell extracts and assayed for 30 min in the absence or presence of 5 µM ADP-heptose (ADPH) or 100 µM UDP-α-D-mannose (UDP-M), ADP-D-ribose (ADP-R) or GDP-α-D-mannose (GDP-M), and the ^32^P-radioactivity incorporated into GST-TIFA quantified (see Methods). **(B)** As in (A), except comparing ALPK1[Ser277Phe] (S277F, pink bars) and ALPK1[Arg150Ala/Ser277Phe] (R150A/S277F, unfilled bars). **(A, B)** The results are expressed as the mean with error bars indicating SEM for two independent experiments, each performed in duplicate.

### The roles of Ser277, Tyr254 and Thr237 in ADP-heptose binding

We compared the location of Ser277, Tyr254 and Thr237 within the crystallographic structure of the ADP-heptose binding domain of ALPK1, which has been solved in the ADP-heptose-bound state (pdb: 5z2c) [1]. Thr237 is situated within the ADP-heptose binding site itself, where it forms a hydrogen bond with a hydroxyl group on the sugar moiety of ADP-heptose (**Fig 4A**). In contrast, Tyr254 is located within an α-helical region just outside the ADP-heptose binding site where, interestingly, it is in close proximity to Ser277 on an adjacent α-helix with which it forms a hydrogen bond (**Fig 4A**). These observations raised the question of whether ALPK1[Ser277Phe] loses its specificity for ADP-heptose and allows activation by human nucleotide sugars due to the loss of the hydrogen bond between Tyr254 and the hydroxyl side chain of Ser277, or due to the mutation of Ser277 to an amino acid with a large hydrophobic side chain (Phe). To investigate this question, we made the Tyr254Phe and Ser277Ala variants to prevent hydrogen bond formation between Tyr254 and Ser277 without affecting the size of these side chains significantly. We found that both of these variants behaved like WT ALPK1 in transfected cells (**Fig 4B**), indicating that it is the bulky hydrophobic side chain introduced by the Ser277Phe mutation that is responsible for modifying the specificity of the ADP-heptose binding site.

**Figure 4:**
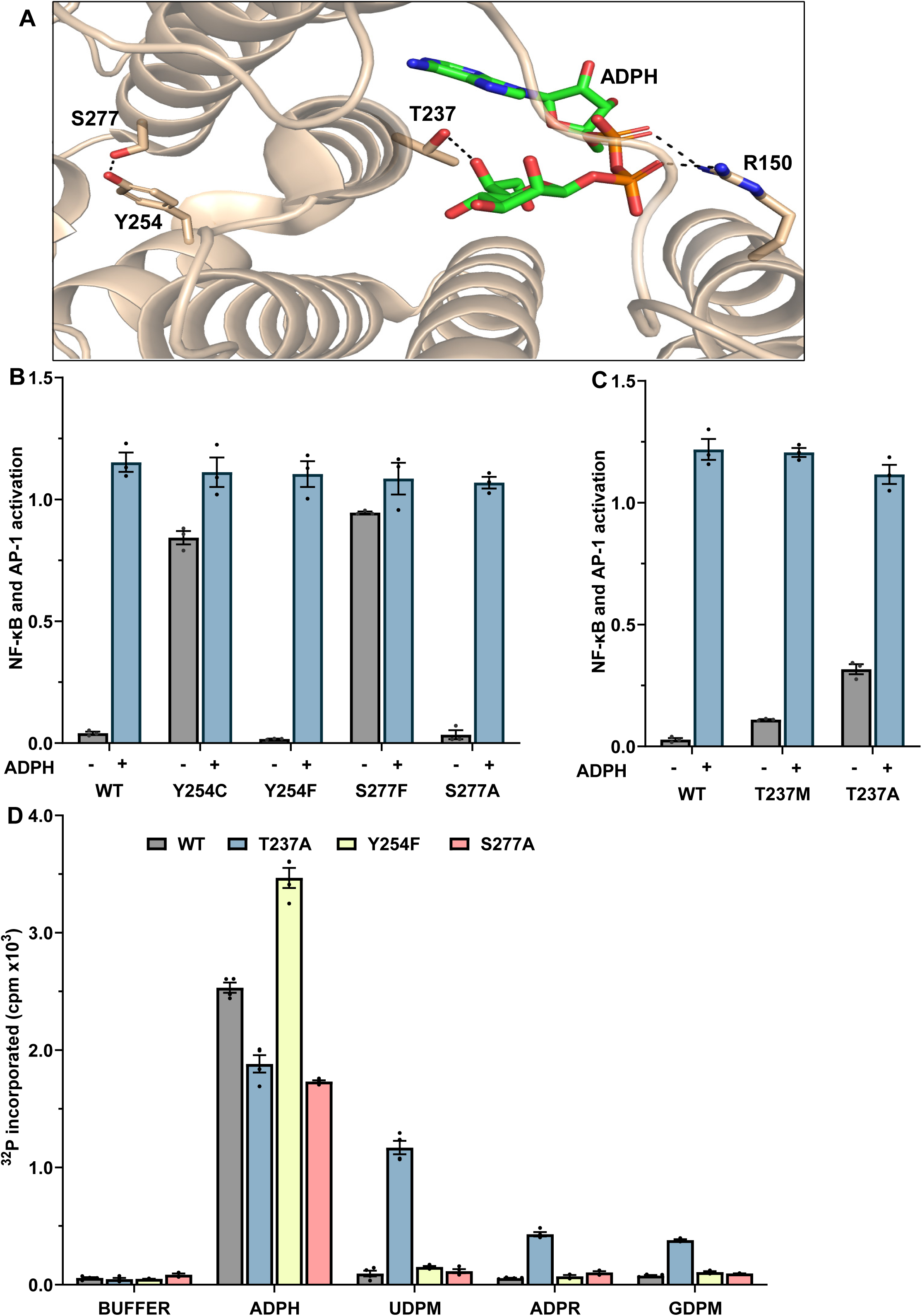
Interactions between Ser277 and Tyr254, and between Thr237 and ADP-heptose, within the ADP-heptose binding domain of ALPK1. **(A)** Location of Ser277, Tyr254, Thr237 and Arg150 in the ADP-heptose (ADPH) binding domain of ALPK1 (pdb: 5z2c). Thr237, Tyr254, Ser277 and Arg150 are shown in stick representation and coloured by element (carbon: bronze; nitrogen: blue; oxygen: red). ADPH is also shown in stick representation and coloured similarly (except carbon: green; phosphate: orange). The interaction between Thr237 and Arg150 with ADP-H, and between Tyr254 and Ser277 are shown by black broken lines. **(B, C)** ALPK1 KO cells were transfected with plasmids encoding WT ALPK1 or the indicated ALPK1 mutants and 24 h later incubated with (blue bars) or without (grey bars) 5 μM ADP-H. The activation of NF-κB/AP-1-dependent gene transcription was then measured after a further 24 h (see Methods). Results are shown as the mean with error bars indicating SEM from an experiment performed in triplicate, with similar results obtained in two additional experiments. **(D)** FLAG-tagged WT ALPK1 (WT, grey bars), ALPK1[Ser277Ala] (S277A, blue bars), ALPK1[Tyr254Phe) (Y254F, yellow bars) and ALPK1[Thr237Ala] (T237A, pink bars) were immunoprecipitated from cell extracts and assayed as in Fig 2. The results are expressed as the mean with error bars indicating SEM for two independent experiments, each performed in duplicate.

The p.Thr237Met mutation disrupts the hydrogen bond formed between the hydroxyl side chain of Thr237 and a hydroxyl group in ADP-heptose. To probe the importance of this interaction in maintaining the integrity of the ADP-heptose binding site, we changed Thr237 to Ala, whose side chain is smaller than methionine. Interestingly, the ALPK1[Thr237Ala] mutant displayed even higher ADP-heptose-independent activity than ALPK1[Thr237Met] in transfected cells (**Fig 4C**). Finally, we carried out cell-free ALPK1 assays on these mutants. Consistent with the cell transfection experiments, the Tyr254Phe and Ser277Ala mutants, like WT ALPK1, were not activated by UDP-α-D-mannose or ADP-D-ribose (**Fig 4D**), while the Thr237Ala mutant, like the Thr237Met mutant, was activated by UDP-α-D-mannose and ADP-D-ribose. Moreover, like the Ser277Phe mutant, but unlike the ALPK1[Thr237Met] mutant, the ALPK1[Thr237Ala] mutant could also be activated by GDP-mannose (**Fig 4D**). These results establish that the hydrogen bond formed between the hydroxyl group of Thr237 and ADP-heptose in WT ALPK1 has an important “gatekeeping” function in preventing interaction with human metabolites. Patients with ROSAH syndrome and a Thr237Ala variant have not yet been identified.

The finding that the *ALPK1* variants causing ROSAH syndrome can be activated by human nucleotide sugars that distinct from ADP-heptose, implies that the conformation of the ADP-heptose binding pocket is altered significantly from WT ALPK1. It should therefore be possible to exploit this difference to develop small molecule drugs that prevent activation of the ALPK1 variants causing ROSAH syndrome without affecting WT ALPK1. Since ROSAH syndrome shows an autosomal dominant inheritance pattern with patients expressing both the wild type and variant forms of ALPK1, such drugs should not impair the bacterial defence mechanism conferred by WT ALPK1 or patient susceptibility to bacterial pathogens.

## Acknowledgements

The project was supported by Wellcome Trust Investigator Award 09380/Z/17/Z (to P.C.)

## Ethical statement

All procedures were performed in accordance with the Helsinki Declaration. Exome and genome sequencing was performed after obtaining informed written consent from all affected individuals or their legal guardians.

## Data, Materials, and Software Availability

All study data has been included in the article.

## Abbreviations

ALPK1: alpha-protein kinase 1
AP-1: activator protein 1
GAPDH: glyceraldehyde 3-phosphate dehydrogenase
DTT: dithiothreitol
GST: glutathione S-transferase
HEK293: human embryonic kidney 293
KO: knockout
NF-κB: nuclear factor kappa-light-chain-enhancer of activated B cells
OTC: optical coherence tomography
RNFL: retinal nerve fibre layer
SDS: sodium dodecyl sulphate
SNV: single nucleotide variants
TIFA: TRAF-interacting protein with forkhead-associated domain
WT: wildtype

## Supplementary Figures

**Supplementary Figure 1.**
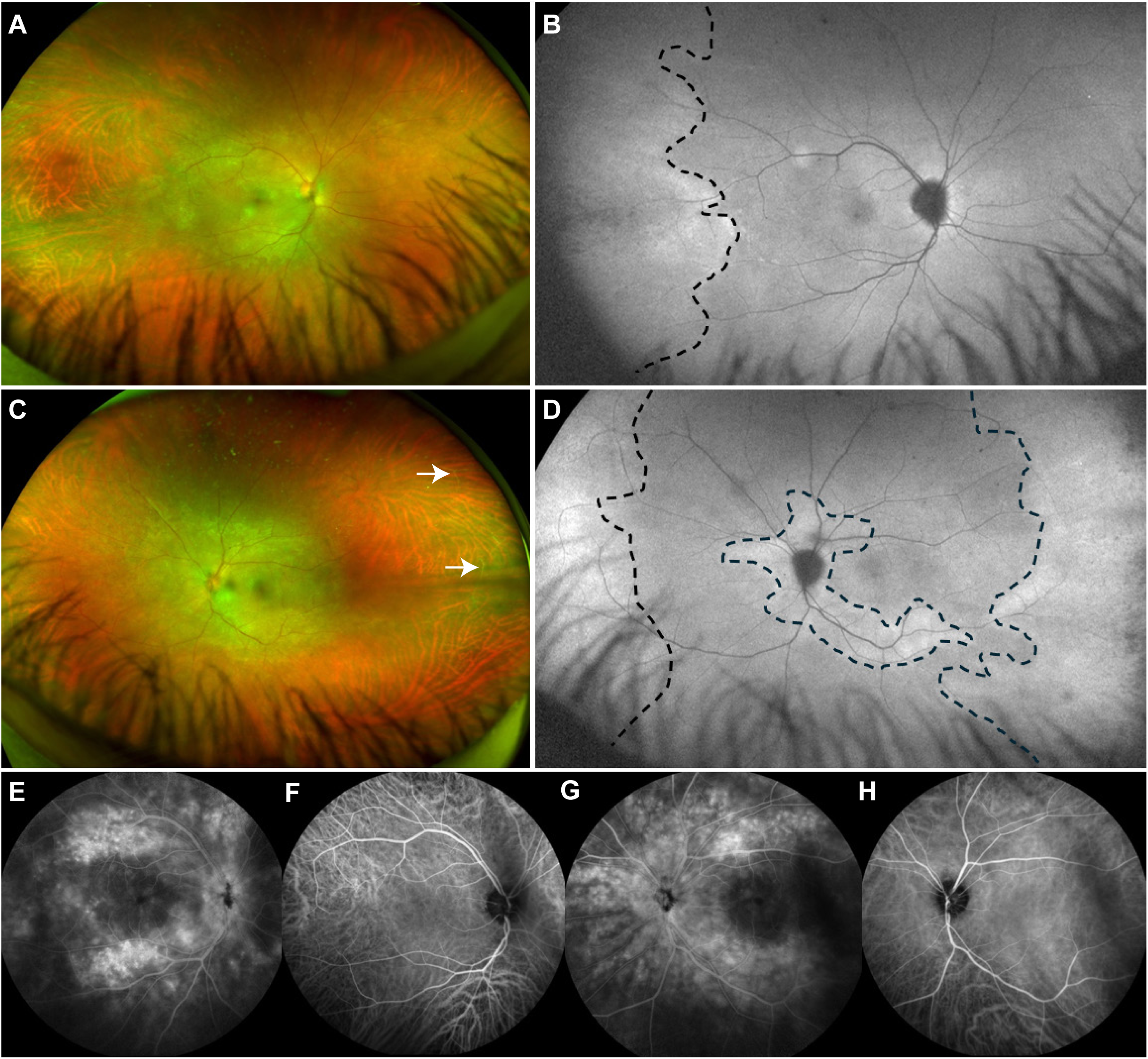
Clinical, angiographic and autofluorescence ophthalmologic findings in Individual II2. **(A,C)** Ultrawide field fundus imaging of right (A) and left (C) eye (Optos, Dunfermline, UK). A mild bilateral optic nerve head oedema is present. Vascular attenuation can be observed in both eyes, predominantly in the temporal side from mid to extreme periphery. Ghost retinal vessels are indicated by arrows. **(B,D)** Ultrawide field fundus autofluorescence (FAF) (Optos, Dunfermline, UK). In the right eye (B), temporal hyperautofluorescence (black dotted line) revealed a photoreceptors/retinal pigment epithelium complex from probable retinal ischemia. In the left eye (D), hyperautofluorescence patterns correspond to the vascular attenuation area seen on fundus imaging and is found in peripapillary zone and infero-temporal vascular arcade, where inflammation is found. **(E-H)** Fluoresceine and Infracyanine Green fundus angiography. Late phase fluoresceine angiography (E,G) shows a large diffusion of juxta-vascular retina near principal vascular arcades and the optic disc in both eyes. Infracyanine Green fundus angiography (F,H) does not exhibit any anomaly during the whole sequence.

**Supplementary Figure 2.**
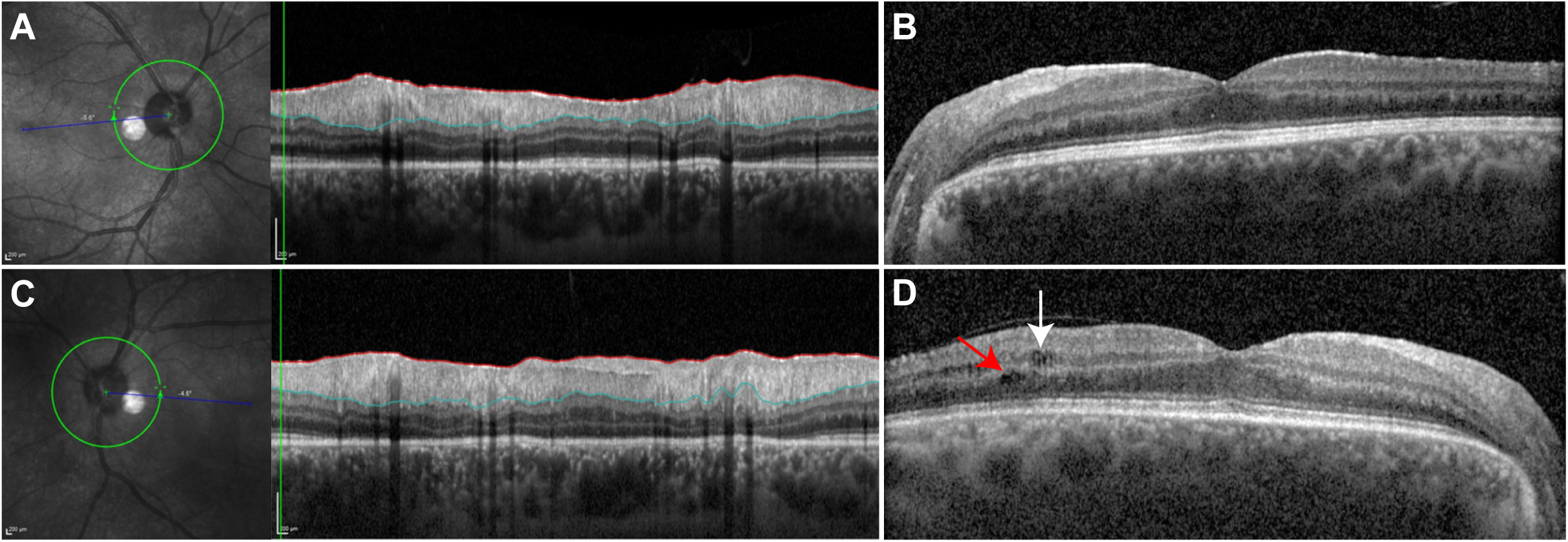
Macular and Retinal nerve fiber layer (RNFL) optical coherence tomography (OCT). **(A,C)** RNFL OCT showing a bilateral thickening of peripapillary nerve fiber layer related to inflammatory oedema. **(B,D)** Macular OCT showing a bilateral retinal thickening, prominent in inner layers. In the temporal side of the right eye (B), intraretinal cysts are present at the level of outer nuclear layer, and in the left eye (D), both in outer nuclear layer (white arrow) and inner nuclear layer (red arrow). In both eyes, the temporal photoreceptor inner/outer segment (IS/OS) junction was seen to be discontinued.

**Supplementary Figure 3:**
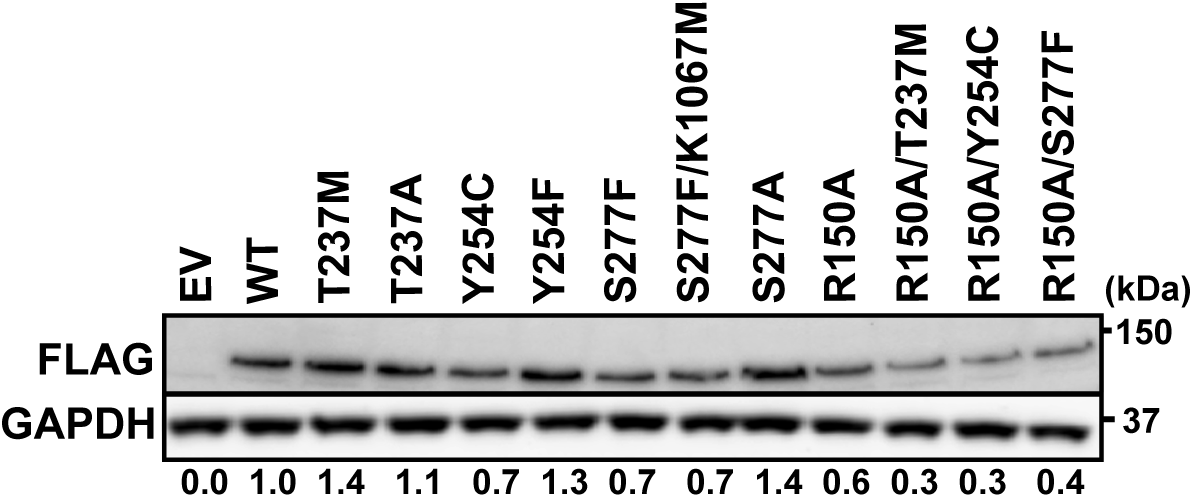
Relative expression levels of different ALPK1 constructs used in this study. ALPK1 KO cells were transfected with empty vector (EV) or plasmids encoding WT ALPK1 or the indicated ALPK1 mutants. After 48 h, cells were lysed, and the extracts subjected to SDS-PAGE and analysed by immunoblotting with the antibodies indicated. The intensities of bands corresponding to different ALPK1 mutants were quantified relative to WT ALPK1 using BioRad Image Lab software (version 6.0.1) and are indicated below the figure.

